# Nanoporous Microelectrodes for Neural Electrophysiology Recordings in Organotypic Culture

**DOI:** 10.1101/2025.05.12.653414

**Authors:** Petro Lutsyk, Debjani Goswami, Stephen Worrall, Stuart D. Greenhill

**Affiliations:** Aston Institute of Photonic Technologies, College of Engineering and Physical Sciences, Aston University, Aston Triangle, B4 7ET, Birmingham, UK; Energy and Bioproducts Research Institute, College of Engineering and Physical Sciences, Aston University, Aston Triangle, B4 7ET, Birmingham, UK; Aston Institute of Health and Neurodevelopment, College of Health and Life Sciences, Aston University, Aston Triangle, B4 7ET, Birmingham, UK

**Keywords:** electrophysiology, organotypic culture, microelectrodes, inkjet printing, aerosol jet printing

## Abstract

Organotypic cultures, specifically brain slices, have been used in neuroscience studies for many years to prolong the lifetime of the biological tissue outside of the host organism. However, the cultures must be kept in a sterile environment, maintaining supply of gas/nutrients for tissue survival and physiological relevance. Electrophysiological recordings from cultured tissue are challenging as the conventional approaches implicate a compromise on biological stability or environmental integrity. In this article, a novel approach has been used to design and print nanoporous microelectrodes on culture wells enabling in situ recording of electrophysiological neural activities. Optimized ink formulations are developed for conductive nanocarbon microelectrodes, and furthermore, fluoropolymer (polytetrafluoroethylene-based AF2400) ink has been inkjet printed for the first time acting as an insulator layer for microelectrodes. To keep the biocompatible nanoporous structure of culture wells, the microelectrodes have been printed on the bottom of the culture cells and only small connector pads have been produced on top of the culture membrane. Neural activity has been recorded by such a microelectrode structure for rodent brain slices cultured for three weeks. Furthermore, aerosol jet printing has been used for printing of nanocarbon microelectrodes allowing to produce much smaller size features compared to the inkjet printing.

## 1. Introduction

Cultured tissues have been used in biological research for many years and are also experiencing a resurgence due to an increasing application of organoids & other multicellular systems. ^[1,2]^ Organotypic cultures of brain slices are utilized by neuroscientists to prolong the lifetime of such tissue outside of the host organism and obtain valuable electrophysiological recordings in vitro, for example, epileptic human tissue obtained through brain resection. ^[3]^ Extracting meaningful functional data from cultured slices is challenging due to the need to compromise the sterile environment of cultures or to remove the culture into a sub-optimal environment to facilitate electrophysiological recordings. Organotypic culture techniques allow preservation of tissue slices in as close to their natural state as possible for weeks, providing an opportunity for long-term experimental manipulation, drug dosing, and other interventions. However, the cultures must be kept in a sterile environment, maintaining proper supply of gas/nutrients for tissue viability.

One of the ways to preserve the culture environment for electrophysiology recordings is to integrate the microelectrodes or microelectrode array (MEA) into culture wells. However, conventional MEAs are fabricated on rigid substrates, and metals are used for electrodes. ^[4]^ The microelectrodes for culture substrate integration need to be biocompatible and porous to replicate the culture well and allow nutrients/gas to access the tissue. Novel materials for microelectrodes have been widely researched, and several studies are confirming that graphene-based MEAs are an excellent candidate for neural recordings due to their biocompatibility, flexibility, and exceptional electrical properties. ^[5,6]^ Viana et al recently demonstrated the feasibility of graphene-based microelectrodes for *in vivo* brain recordings with high fidelity and superior implantation biocompatibility. ^[7]^ The electrode material is recognized as a critical factor that determines the performance of neural interfaces in such implants. ^[7,8]^ Furthermore, neural cell cultures on graphene substrates showed good viability with enhanced neurite growth. ^[9]^ Another outstanding feature of graphene is its ability to form structures with high double layer capacitance at the interface of conductive microelectrodes with electrolyte (i.e., aqueous media). ^[7,8]^ Porous graphene electrodes (produced by various techniques, e.g. EGNITE, ^[7]^ free-standing graphene-fiber, ^[10]^ laser-induced graphene (LIG) directly grown on polyimide ^[11,12]^ or photoresist ^[13]^ substrate using laser pyrolysis, a mixture of graphene and carbon nanotubes ^[14]^) have been explored aiming to overcome the challenging trade-off between porosity and electrode performance. Perforated MEA is an alternative technology for a few days long neural activity monitoring requiring continuous perfusion of media. ^[15]^ To the best of our knowledge, there are no studies of integrating media-permeable nanoporous microelectrodes to culture well substrates, enabling continuous monitoring of neural activity of organotypic cultures.

In this paper, the innovative approach has been used to design and print microelectrodes on highly porous culture wells allowing to record electrophysiological neural activity for rodent brain slices. To keep the biocompatible nanoporous structure of culture wells, the microelectrodes were double side printed on the culture substrates with insulative layers on the bottom and only a small connector pad contacting cultured tissue on the top (**Figure 1**).

**Figure 1.**
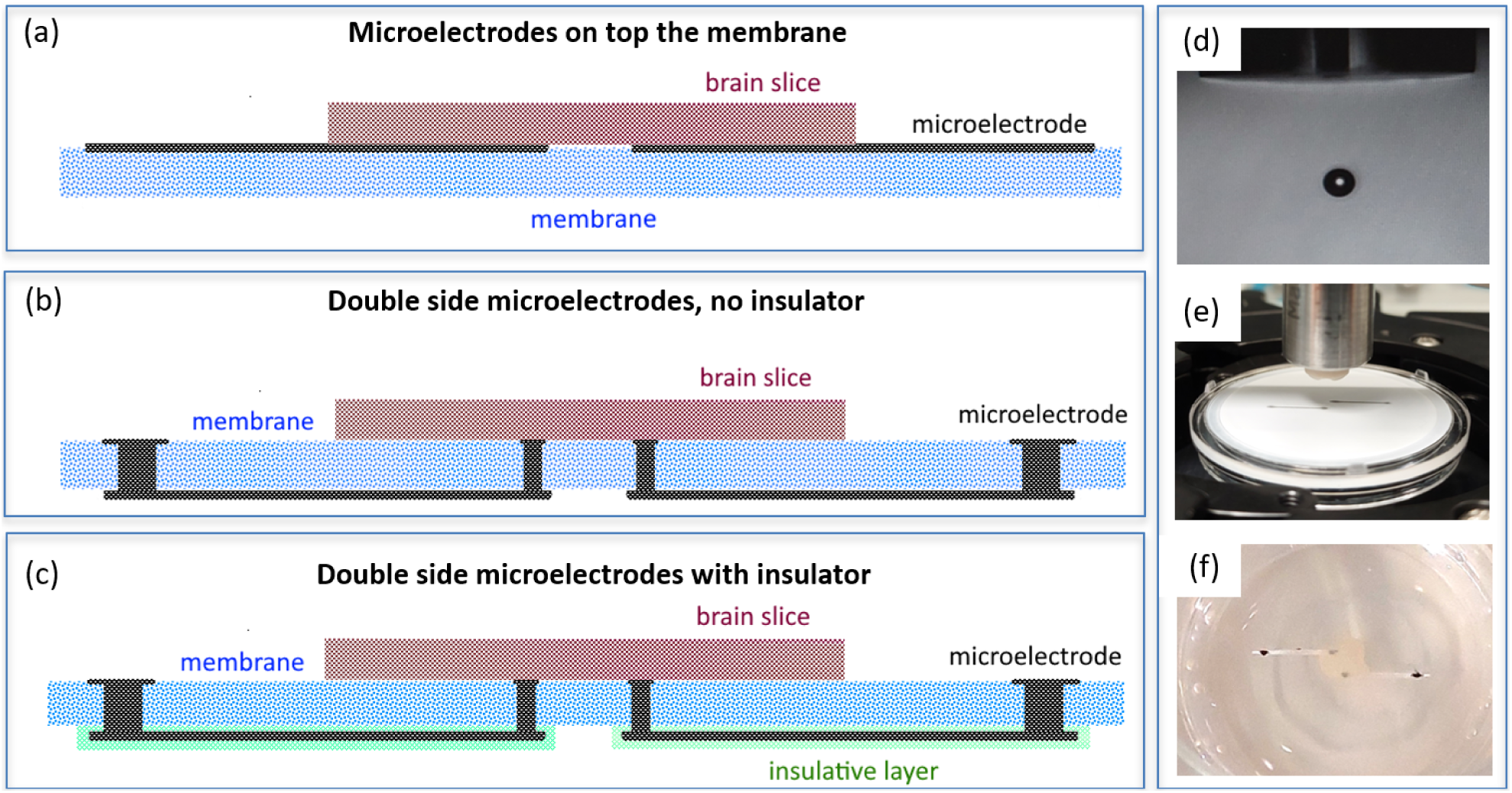
(a-c) Schematic architecture of the microelectrodes on the membrane for organotypic culture, where (a) microelectrodes were printed only on top of the membrane, (b) double side microelectrodes (b) without and (c) with insulator at the bottom of the membrane; (d) nanocarbon ink drop produced from 50 µm nozzle; (e) inkjet printing setup with 30mm Millicell culture plate insert; (f) cultured brain tissue in the culture well with microelectrodes printed as in the part (c).

Designed microelectrode structure enabled neural recordings for the tissues cultured for three weeks. Optimized ink formulations were developed for conductive nanocarbon microelectrodes and insulative polymer layers. Importantly, inkjet printing of PTFE-based polymer layers has been demonstrated for the first time. The presence of the insulative layer for microelectrodes was a key factor in obtaining good quality recordings, as the microelectrodes without insulative layers (Figure 1a,b) had substantial dissipation of the electrophysiology signal to the fluids preventing to record electrical activities of organotypic cultures.

## 2. Results and Discussions

### 2.1. Conductive Ink Formulation

Conductive nanocarbon microelectrodes have been produced by application of shear-force mixing with 1-pyrenesulfonic acid (PS1) as surfactant. Pyrene sulphonic derivatives are proven to be one of the most efficient surfactants in producing water-based stable (up to 1 year) dispersions of graphene and graphene-based nanocarbon. ^[16,17]^ Such dispersions yield high concentrations of nanocarbon without oxygen groups ^[16]^ and are shown to be inkjet printable ^[17]^ even considering that their inverse Ohnesorge number *Z* is higher than 14. *Z* is expected to be in the range of 1 < *Z* < 14 to produce stable drops. ^[18]^ For inkjet printer settings with nozzle diameter (*d*) = 50 µm, water-based ink would produce *Z* ∼ 60, much exceeding the above range. Therefore, additional refinements for nanocarbon ink formulations were required. Adding Triton X-100 allowed us to reduce surface tension from 72.8 mN m^-1^ (water) to about 33 mN m^-1^. Propylene glycol admixed to our printing solution resulted in the viscosity increasing up to 2 mPa·s. Due to the reduced surface tension and increased viscosity, our nanocarbon ink has *Z* ∼ 20 for a nozzle diameter of 50 µm (Table S1). Such ink produced a stable drop (Figure 1d) even with *Z* being slightly above 14; a similar outcome was shown before. ^[17]^ Furthermore, the resulting ink had diminished coffee-ring effect (due to weaker Marangoni flow ^[19]^). Finally, the obtained nanocarbon ink is suitable for aerosol jet printing via ultrasonic atomization, which works well for inks with a maximum viscosity of 5 mPa·s.[20]

It was previously shown that shear-force mixing is a scalable, defect-free, and highly efficient technique for graphene exfoliation (alternative to sonication) and, as a proof-of-concept, these studies used sodium cholate as a surfactant to disperse graphene in water. ^[21,22]^ Application of PS1 for graphene exfoliation required 72 hours of tip sonication to produce highly concentrated PS1-based ink. ^[17]^ We have applied shear-force mixing with PS1 to produce nanocarbon ink, and only 2 hours of treatment was needed to reach the required level of nanocarbon concentration.

### 2.2. Insulative Ink Formulation

For insulative ink, we have used perfluorodecalin solution of AF2400 (5 g L^-1^). The polymer was chosen because of its excellent gas permeability due to the inherent extensive free volume ^[23,24]^ and having a molecular structure similar to the culture membrane substrate (made of PTFE). Surface tension and viscosity of such solution were 27 mN m^-1^ and 5.5 mPa·s, respectively (slightly increasing compared to the features of neat perfluorodecalin). This ink formulation has *Z* = 9.5 for *d* = 50 µm (Table S1), and a stable drop was formed during inkjet printing. Apart from having *Z* within the expected range, polymer-ink printing needs to follow viscoelastic jet stability to avoid ‘bead-on-a-string’ structure achieved for flexible polymers of sufficiently high molecular weight. ^[25]^ Therefore, the solution of AF2400 has been tip sonicated, aiming to reduce the molecular weight of the polymer and ensure more stable drop formation. Ultrasonication is proven to reduce polymer chain length, lower the viscosity of the ink, and allow for higher polymer concentrations and improved printability.^[26]^ An alternative way to increase the concentration of polymer is via application of an oil-in-water emulsion, ^[27]^ however, this technique requires extra additives that might affect the biocompatibility of microelectrodes. A less viscous solvent (co-solvent) or an admixture of surfactant to lower the surface tension of the resultant ink could be applied for printing of AF2400 in case higher concentrations of the polymer ink are required for alternative applications. The AF2400 concentration has been optimized towards high-level values for perfluorodecalin-based ink, resulting in stable drops without side droplets (which is also a common issue in polymer inkjet printing ^[28]^). Drop formation stability and dispenser head nozzle clogging issues have been observed for the AF2400 concentrations above 5 g L^-1^.

### 2.3. Ink Characterization by Absorption, Raman, and Photoluminescent Spectroscopy

The nanocarbon concentration in the inks was determined using absorption spectroscopy in the visible range. The absorption spectrum of graphene and larger graphite nanoparticles is nearly flat and has no features in the long-wavelength visible region (**Figure 2**a). There are extensive studies in which the absorption is measured at 660 nm for the concentration estimation using the Beer–Lambert law. ^[21,22]^ The concentration estimation by absorption coefficient for such material has some variance, ^[29]^ therefore, in this study a conventional value for absorption coefficient at 660 nm (*α*_660_ = 2460 L g^-1^ m^-1^) ^[30]^ was used. By this technique, nanocarbon concentration in the ink was evaluated to be in the range of 1 g L^-1^. The nanocarbon ink still has some content of PS1, as the characteristic PS1 absorption peak at 375 nm is present in the ink spectrum (Figure 2a). Comparing the intensity of the 375-nm peak in the ink and the neat PS1 solution (Figure 2a), the concentration of PS1 in the ink is estimated to be ca. 0.3 g L^-1^. Insulative ink, perfluorodecalin solution of AF2400, has the absorption maximum of 206 nm and is practically transparent in the UV, visible, and near infrared range from 250 nm (Figure S1).

**Figure 2.**
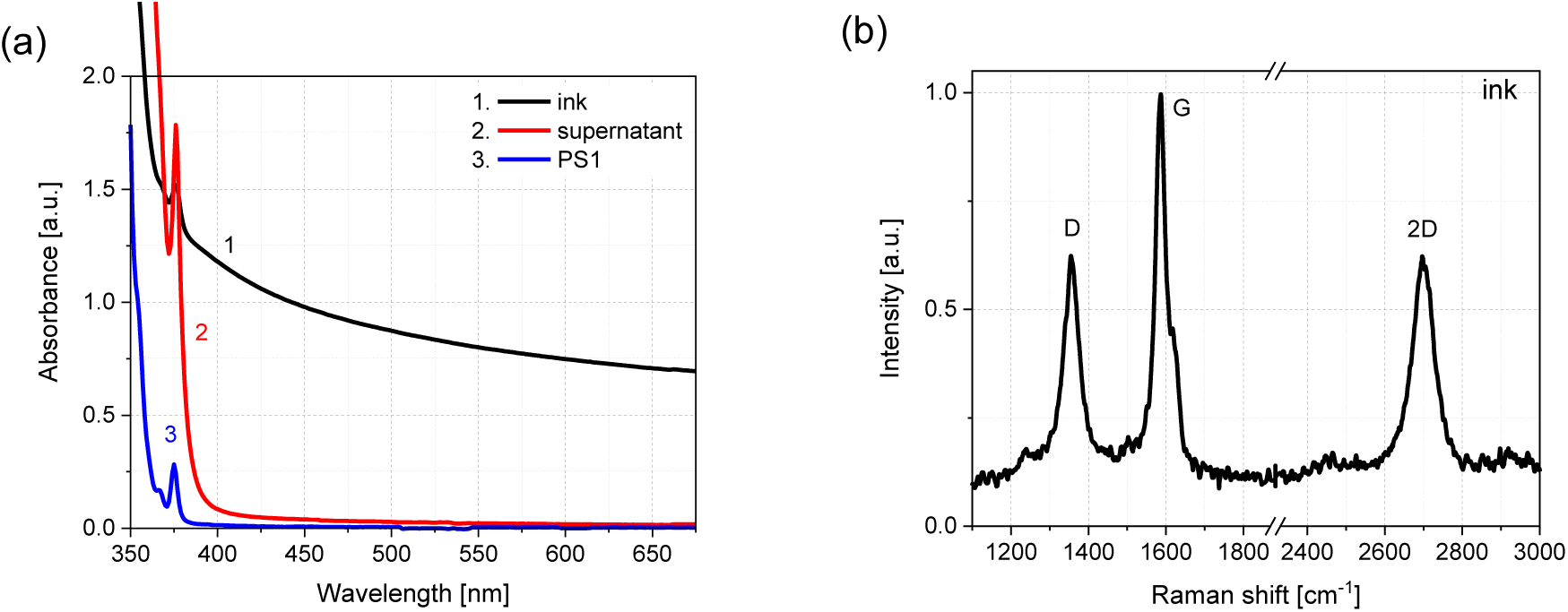
(a) Absorption spectra of nanocarbon supernatant obtained in the process of ink production (curve 1; diluted 1:6), the nanocarbon ink (curve 2; diluted 1:9) and PS1 at the concentration 0.1 g L^-1^. All measured in 3mm quartz cell. (b) Raman spectra of the nanocarbon ink deposited on SiO_2_ substrate after washing in water (to remove excess PS1).

Raman spectra of the ink deposited on SiO_2_ (with PS1 removed by washing in water) exhibit characteristic carbon peaks at 1354 (D), 1587.5 (G), and 2696 (2D) cm^-1^ (Figure 2b). Such spectral features, particularly the 2D peak, indicate the presence of (2-4)-layer graphene nanoparticles and no graphitic flakes, thus evidencing efficient exfoliation.^[31]^ Relatively wide G and 2D bands, as well as the high D/G ratio, are indicative of substantive defect/edge density and small lateral dimensions of the obtained nanocarbon ink nanoparticles.^[32]^ Photoluminescence (PL) measurements demonstrated that the nanocarbon ink is featured by monomer emission of PS1 molecules in the range of 370-430 nm (**Figure 3**). Such monomer emission has characteristic narrow peaks at 374, 393, 413 nm and small spectral bands at 382 and 403 nm. However, the most significant PL feature of the ink is a lack of broad excimer emission of the PS1 in the range of 430-600 nm (peaking at 494 nm) that is present in the highly concentrated aqueous solution of PS1 (Figure 3c, curve 3) and supernatant obtained during the ink fabrication (Figure 3c, curve 2). The excimer emission evidences the formation of dimers of PS1 molecules and demonstrates a substantial excess of PS1 surfactant in the supernatant.

**Figure 3.**
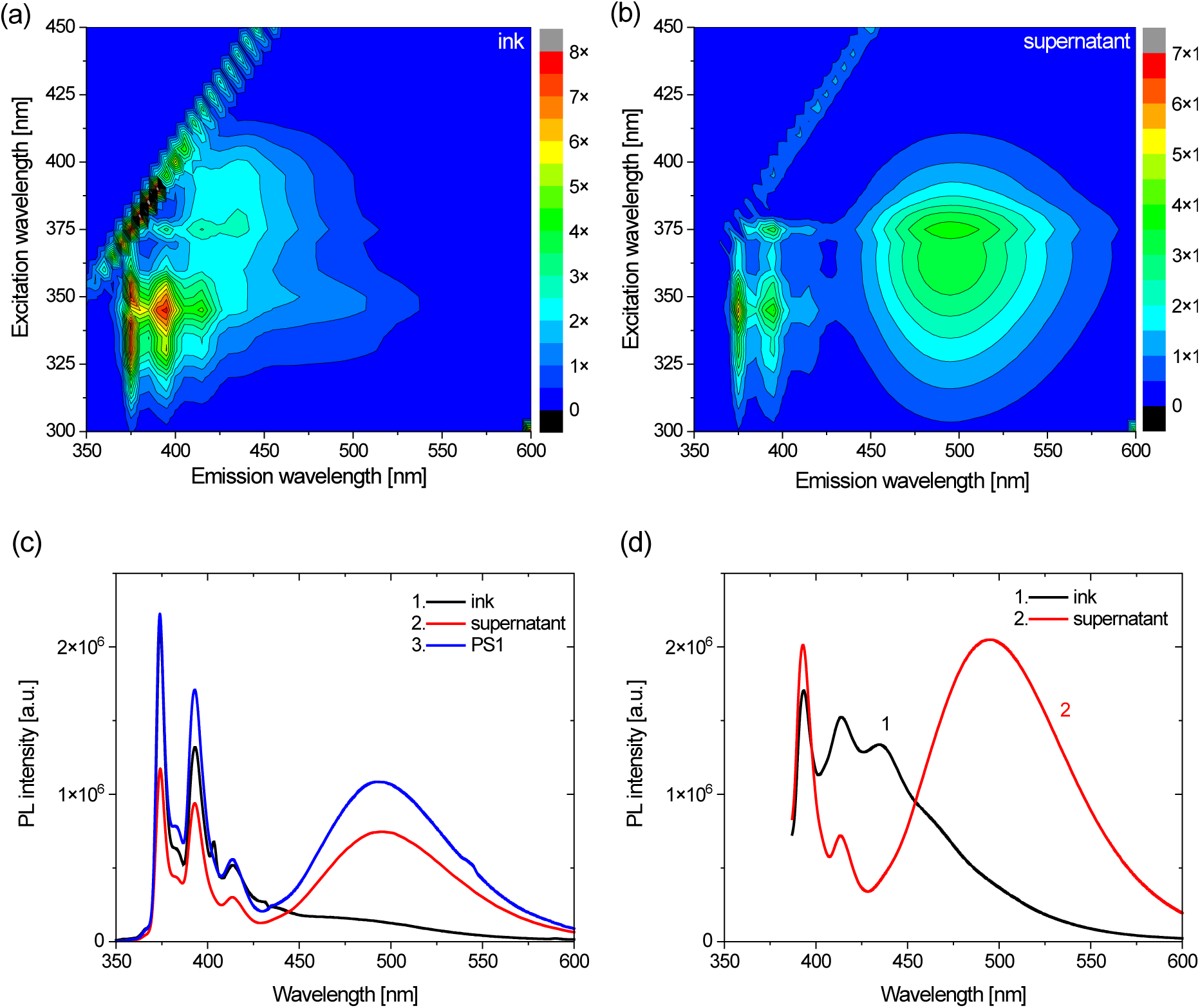
(a,b) Excitation-emission photoluminescence (PL) maps of nanocarbon ink (a) and supernatant obtained in the process of ink formulation. (c,d) PL spectra of nanocarbon ink (c,d - curves 1) supernatant obtained in the process of ink formulation (c,d – curves 2); and aqueous solution of PS1 at the concentration of 6 g L^-1^ used in the ink formulation (c – curve 3); excitation wavelengths are 310 (c) and 375 (d) nm.

Another feature of nanocarbon ink emission is an amplification of PL band at 413 nm and the emergence of a new band at 434 nm and a broad shoulder in the range of 450-475 nm. These ink features are appearing at excitation wavelength range 340-400 nm (Figure 3a) and can be associated with weak emission of nanocarbon particles in the dispersion.

### 2.4. Inkjet and Aerosol Jet Printed Microelectrodes

The hydrophilic PTFE membrane of Millipore culture inserts with 0.4 µm pores (according to the specs) has a highly branched network of polymer micro-sized wires connected to multiple nodes, and this is well evidenced by our SEM measurements (**Figure 4**a at 5 µm scale and Figure S2 at 2 and 5 µm scales). Surface porosity of neat PTFE membrane obtained from SEM imaging is in the range of 63-66%, and together with its high-free-volume internal mesh structure should be responsible for high viability (for as long as 40 days) and excellent trans- membrane oxygen transport. ^[33]^ Therefore, the porous features of such membranes need to be preserved during the microelectrode’s integration to such substrates.

**Figure 4.**
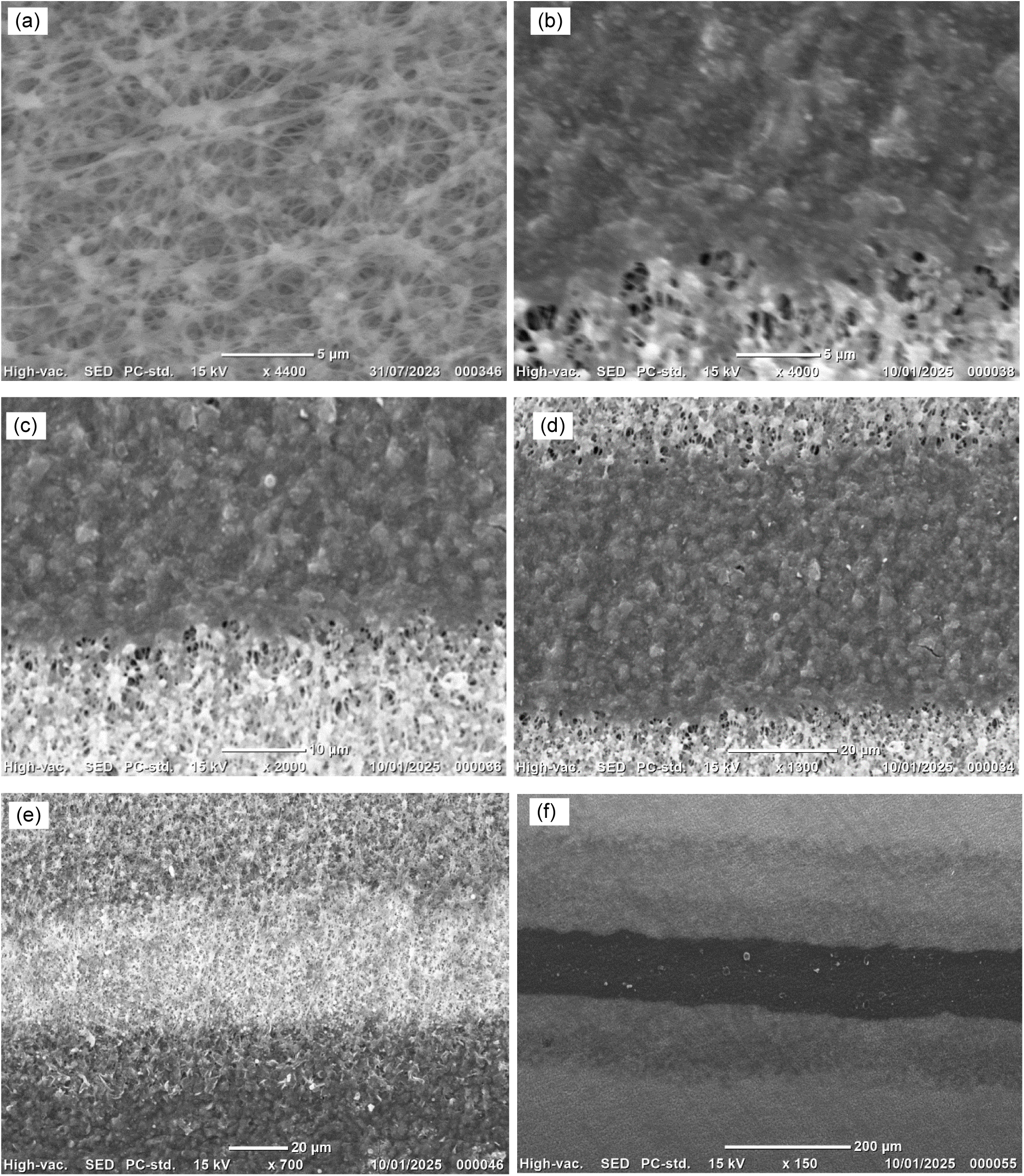
SEM images for (a) neat PTFE membrane of culture insert substrate, (b-d) aerosol jet printed nanocarbon microelectrode on the PTFE membrane; (e) aerosol jet printed microelectrode (bottom) with AF2400 insulative cover (middle) on the PTFE membrane (top); (f) inkjet printed microelectrode (middle dark area) with AF2400 insulative cover on the PTFE membrane.

Aerosol jet and inkjet printing of the formulated in this study inks resulted in good quality microelectrode/insulator lines (Figure 4b-e). The width of the aerosol jet printed microelectrode (20 printing passes) is approx. 45 µm, whereas inkjet printed microelectrode width (10 printing passes) extends up to approx. 100 µm. Thus, aerosol jet printing for nanocarbon microelectrodes allowed us to produce much smaller size features compared to the inkjet printing. Besides, the edges of the aerosol jet printed microelectrodes (Figure 4d) are smoother than the inkjet printed lines (Figure 4f) due to a stable aerosol jet with little or no overspray (minimizing overspray is identified as one of the major challenges in the initial process development for aerosol jet printing ^[20]^).

Inkjet printing of AF2400 ink resulted in the formation of widened insulation lines of approx. 200 µm (for 20 printing passes). Wider lines of AF2400 ink are formed due to a high wettability of perfluorodecalin-based ink with the PTFE membrane (Figure S3-S5). Inkjet printing of insulator lines ‘sandwiching’ conductive lines works very well for aerosol jet printing of microelectrodes, allowing ample offset to cover a 45-µm-width of the nanocarbon conductor by a 200-µm-width AF2400 insulator. Initial biocompatibility studies showed that a high number (>40) of printing passes for insulative ink resulted in a filling of the PTFE nanopores and the membranes losing permeability feature. Therefore, too much of AF2400 insulative cover either on top or at the bottom of the PTFE membrane significantly reduces the viability of the organotypic cultures due to reduced trans-membrane porosity and subsequent supply of media supplements/O_2_/CO_2_. Aiming to achieve reasonable insulation but preserving the nanoporous structure of the culture insert substrate, AF2400 insulative cover required thickness optimization, where having 20 printing passes provided good outcomes with a trade-off between biocompatibility and insulation. SEM images (Figure 4e) demonstrate that the areas of AF2400 insulative cover still have pores, but the branched network has wider PTFE membrane wires being covered by AF2400. Surface porosity (obtained from SEM images) of such optimized AF2400 layers was in the range of 46-50%, whereas nanocarbon ink porosity ranged within 39-43%, evidencing some reduction in the porous area on the PTFE surface. AF2400 and nanocarbon prints have an excellent adhesion to the PTFE membranes, as well as nanocarbon prints on PTFE covered by AF2400.

The microelectrodes were characterized with impedance spectroscopy (**Figure 5**). Impedance at 1kHz is ca. 0.9 MΩ. Such values are in the range of previously reported impedance for nanocarbon microelectrodes ^[6,34]^ and are below the 3MΩ-benchmark, impedances above which potentially leading to increased noise and reduced signal-to-noise ratio in electrophysiology recordings. ^[35]^ Flattening of low-frequency impedance for nanocarbon- based microelectrodes has been evidenced in literature ^[34,36]^ featuring resistance of the microelectrodes. For frequencies higher than 1kHz, capacitive behavior starts to dominate reducing the impedance below 10 kΩ in the range of 1MHz (Figure 5). Flattened shape of the impedance spectrum below 1kHz is a good aspect for neural signals recordings, where most signals have a frequency in this spectral range and would have minimal impact of impedance variations. Impedance spectra have been modelled by a parallel combination of the Constant Phase Element (CPE; representing capacitive behavior) and charge transfer resistance (*R*_ct_), shown in Figure 5b. The fitting parameters are summarized in Table S2, allowing the evaluation of double layer capacitance ^[37]^ and effective surface area of the microelectrodes being in the range of (0.10 - 0.25) nF and 10^-5^ cm^2^, respectively. The formulas are added to the Supporting Information file. Furthermore, the frequency spectrum of noise (Figure S6) is practically flat in the range up to 1kHz (50 Hz spike represents power line hum), evidencing no impact of shot noise or other frequency-dependent artefacts.

**Figure 5.**
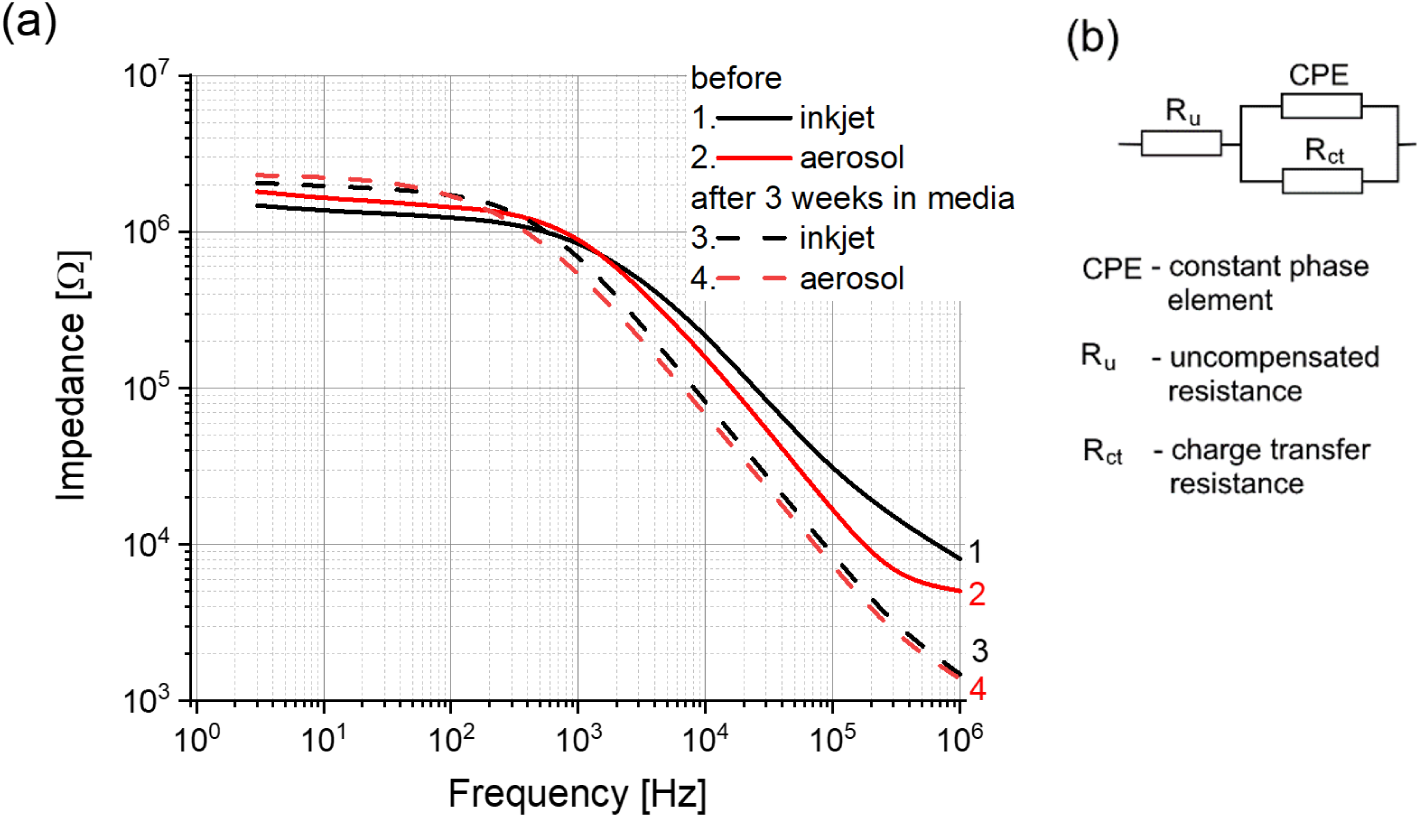
(a) Impedance spectroscopy of inkjet (1,3) and aerosol jet (2,4) printed nanocarbon microelectrodes before (1,2) and after keeping the system 3 weeks in the media environment (3,4). Inkjet and aerosol jet printing had 10 and 20 passes, respectively. (b) Scheme of equivalent circuit by the Constant Phase Element (CPE) model used to fit the impedance spectra.

It needs to be noted that the conductivity/impedance of the microelectrodes could be improved by thermal treatment, but such step has been eliminated at this stage of the studies to avoid additional impact on the biocompatibility of Millipore culture insert membranes.

Furthermore, nanocarbon ink formulation can be further optimized, or an alternative treatment of microelectrodes applied to improve the electrical properties of printed lines.

Subtle changes of impedance were observed after keeping the microelectrodes in the media for 3 weeks (Figure 5), and, as result of such prolonged storage in the media, the shapes of impedance spectra became nearly same for of inkjet (1,3) and aerosol jet (2,4) printed microelectrodes (Figure 5, curves 3,4). There is a little increase of the impedance in the low-frequency range (below 100 Hz) and moderate decrease of impedance in the high-frequency range (above 1kHz) due to the media (Figure 5). The decreased impedance indicates a reduction in capacitive reactance for the microelectrodes. Such behavior can be attributed to the adsorption/desorption of specific ions from nanoporous microelectrodes.^[37]^ Moreover, such prolonged keeping of the microelectrodes in the media environment did not result in any noticeable deterioration of the microelectrode physical appearance. Overall, this evidences a feasibility of the technology for producing stable conductive microelectrodes/insulative layers on the culture substrates.

The initial design of microelectrodes was to print all the structures on top of the membrane (Figure 1a). However, the signal from such microelectrodes without insulative layer was very low, whereas adding even very thin insulation on top of the membrane resulted in a compromised biocompatibility. Therefore, to maintain the nanoporous structure of culture wells biocompatible, double side printing of microelectrodes has been considered (Figure 1b), where most of the printing is happening at the bottom and only small connector pads are designed to be on top of the substrate. The double side printing without insulative layer covering microelectrodes (Figure 1b), same way as schematics on top of the membrane only (Figure 1a), resulted in substantial dissipation of the electrophysiology signal to the conductive media of organotypic cultures inhibiting registration of the electrical pulses. Thus, the double side microelectrodes with insulative layer (at the bottom of the substrate) covering most of the conductive parts have been designed (Figure 1c). Such schematics leave uncovered only a small connector pad on top of the culture membrane that connects to the brain slice and another connector pad on the opposite side of the microelectrode that is used for connection to the signal registration system. The microelectrodes, as shown in Figure 1c, appeared to be highly efficient in recording electrophysiology signals with a high signal-to-noise ratio and good biocompatibility.

PL studies for printed microelectrodes allowed us to understand additional features of such systems. As-printed microelectrodes have distinctive emission peaks at 375, 393, and 413 nm with excitation maximum at 345 nm (**Figure 6**a). These intense peaks evidence a significant presence of PS1 molecules on the substrate with as-printed lines. It needs to be noted that PS1 is spreading on the PTFE membrane due to the wetting process, particularly for multiple printing passes. For nanocarbon ink, the carbon particles (a few hundred nm in size) stay in the place of the ink drop landing on the surface, whereas water (and consequently water-soluble surfactant molecules) can penetrate the hydrophilic PTFE membrane and spread further. Such a process becomes visible with the naked eye after several printing passes made in a very short time, as the membrane around the printed microelectrode changes color (from non-transparent white to pale translucent). As water gradually evaporates, the membrane around the microelectrode returns to non-transparent white with a slight staining of grey.

**Figure 6.**
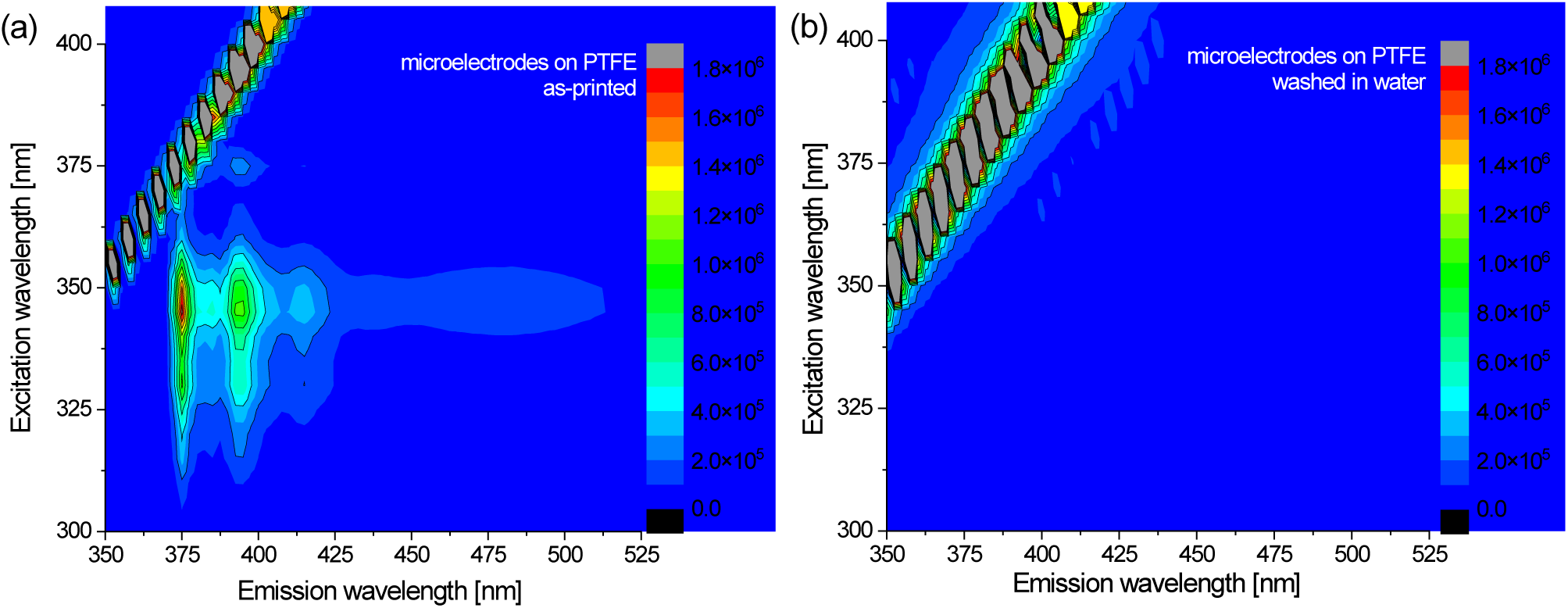
Excitation-emission photoluminescence (PL) maps of the microelectrodes (a) as printed and (b) after washing in water (to remove excess PS1).

PS1 is well soluble in water, and the follow-up washing of the microelectrodes in water has resulted in disappearance of PS1 peaks (Figure 6). It evidences efficient removal of the PS1 molecules from printed microelectrodes. Removing PS1 from microelectrodes allowed us to observe a low-intensity broad PL band having a maximum in the range of excitation wavelengths 380-400 nm and emission wavelength 450-600 nm that features well a neat PTFE membrane of the Millicell culture plate inserts (Figure S6). In this broad band, there is a potential contribution of the low-intensity PL shoulder from nanocarbon, evidenced in Figure 3a (PL emission in the range of 450-475 nm at excitation wavelength range 340-400 nm). Low intensity broad emission from nanocarbon microelectrodes (on PTFE membrane) provides a good background for potential *in situ* PL studies of organotypic cultures (e.g., simultaneous optical imaging and electrophysiological recording ^[34]^).

### 2.5. Electrophysiology Recordings and Histology

After 21 days in culture using both insulated and non-insulated nanocarbon-printed culture wells, electrophysiological and histological analysis were carried out on the brain slices to assess cellular and network function, and to confirm biocompatibility of the printed electrodes. To provoke widespread activity, the GABA_A_ antagonist gabazine (20µᴍ) was added to the culture wells in order to disinhibit the slice.

As would be expected, the clearest and most reliable recordings were obtained from wells, which had insulating polymer encapsulating the printed nanocarbon traces (**Figure 7**a,b). With exposed nanocarbon traces, short-circuiting of the neuronal signals into the bath aCSF, and too-low impedances reduced the quality and reliability of the recordings. With insulated traces, cultured slices were able to be aligned with the recording pads to ensure that identifiable anatomical landmarks (e.g., CA1 of the hippocampus) were located on top of the recording site. Placing either high-impedance (Figure 7a) glass or low-impedance solid wire (Figure 7b) electrodes on the relevant recording pad resulted in a multi-unit recording of neuronal action potentials that was not evident when the electrode was not directly making contact with the pad. Root mean square (RMS) noise (calculated from post-mortem recordings using inkjet printed microelectrodes) was 0.011±0.001 mV and 0.26±0.01 mV for the glass and solid wire connections, respectively. The average signal-to-noise ratio (calculated from the ratio of average peak-to-peak spike signals divided by twice the RMS noise voltage: the peak-to-peak signal average/(2 × RMS noise)) was in the range of 12. RMS noise of aerosol printed microelectrodes was around two times lower, resulting in improved signal-to-noise ratio (ca. 24).

**Figure 7.**
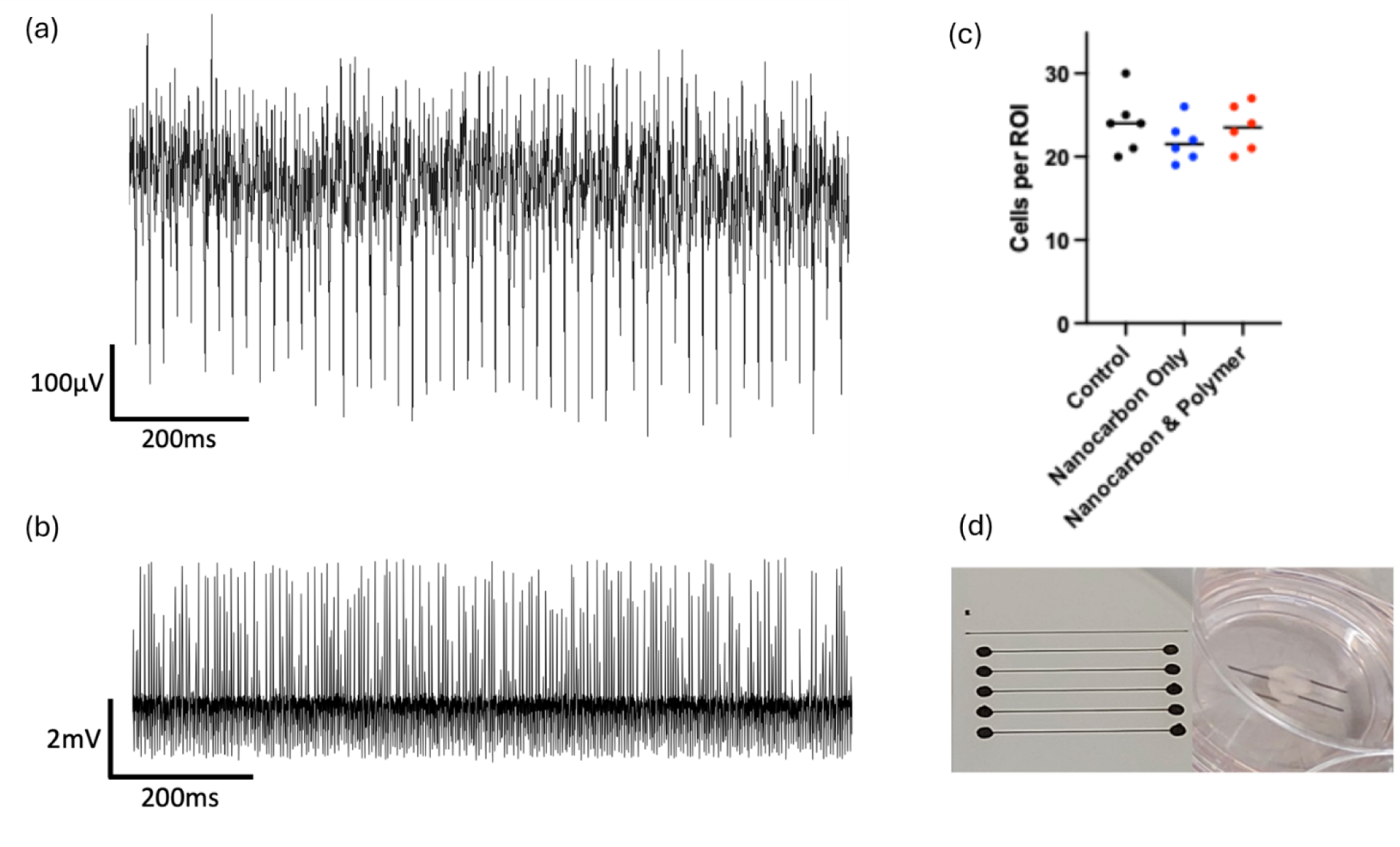
Electrophysiology and biocompatibility of novel culture system. (a) High-impedance glass microelectrode recordings show well-defined multi-unit activity in a cultured brain slice in the presence of gabazine. (b) Low-impedance wire electrode recordings also show multi-unit activity with some rebound shot noise. (c) Cell counts show no significant difference in histology between control, nanocarbon and insulated nanocarbon conditions. (d) Example pad configuration (left) and brain slice in situ (right).

Cultured slices were also fixed and stained with hematoxylin and eosin to allow for quantification of cell density (Figure 7c). In a 100×100µm region of interest (ROI), there were no significant differences between control cultures, exposed nanocarbon and insulated nanocarbon (*n* = 6 per group, Kruksal-Wallis test; averaged cell counts per ROI were 24.0, 21.5, and 23.5 for control sample, nanocarbon as in Figure 1a and double-printed nanocarbon with insulator as shown in Figure 1c, respectively) suggesting that any cell loss was consistent across all the culture well construction, with no biocompatibility concerns from the printed additions (Figure 7c).

## 3. Conclusions

Novel microelectrodes printed on highly porous culture substrates enabled the recording of multi-unit neural activity of rodent brain slices cultured for three weeks. The high-quality recordings and biocompatible structure of culture membranes were obtained by double side printing of microelectrodes with the addition of insulative layers. Besides, aerosol jet printing of small-sized microelectrodes ‘sandwiched’ by inkjet jet printing of insulator lines resulted in the structures with substantially improved signal-to-noise ratio of electrophysiological recordings. Designing nanoporous microelectrodes on culture wells can pave the way for the development of more advanced setups for long-term monitoring of cultured tissues, enabling transformative studies of brain activities & potential for novel treatment of brain pathologies, including personalized medicine. Besides, this technology can drive the reduction in animal use by increasing the usability of tissue taken from animals. Finally, the feasibility of inkjet printing for PTFE-based polymer has been demonstrated, and this particular outcome can be translated into various applications beyond insulating layers for microelectrodes, for example, tailorable optoelectronics encapsulation structures, optical and microwave windows.

## 4. Experimental Section / Methods

*Conductive Ink Preparation*. Graphite powder (4 g; <20 μm initial size of powder particles; Sigma Aldrich / Merck) was dispersed in the presence of a preliminary dissolved 1- pyrenesulfonic acid (PS1) (240 mg) in deionized water (40 mL). An L4RT high shear mixer (Silverson) was used for dispersion with an emulsion screen at 10,200 rpm and at room temperature sample cooling bath. The obtained dispersion was centrifuged with a Mini Spin plus centrifuge (Eppendorf SE) at 3,700 rpm for 20 min, and the top 50% of supernatant was collected and centrifuged again at 14,200 rpm for 1 hour. Approximately 0.5 mL of precipitate was collected and mixed with 2 mL of a printing solution to obtain the nanocarbon ink. The printing solution was obtained by mixing 2 mL of propylene glycol with 10 mL of deionized water and ≥1 μL of Triton X-100 (all Sigma Aldrich / Merck). The nanocarbon ink was filtered using a 5-µm-pore PTFE syringe filter.

*Insulative Ink Preparation.* Poly[4,5-difluoro-2,2-bis(trifluoromethyl)-1,3-dioxole-co- tetrafluoroethylene] or short AF2400 was dissolved in perfluorodecalin (both Sigma Aldrich / Merck) with the concentration of 5 g L^-1^. The solution was ultrasonicated for 20 min and filtered via a 5-µm-pore PTFE syringe filter to ensure polymer-based ink forms good quality drops for inkjet printing.

*Ink Characterization*. The surface tension and contact angle of the ink were obtained with a L2004A1 Ossila contact angle goniometer, whereas the viscosity was measured via a falling ball viscometer. The absorption spectra were measured with a double beam Lambda 1050 UV/VIS/NIR spectrometer (Perkin Elmer) using 3mm or 10mm quartz cuvettes. Raman spectra were measured with dual-channel automated Raman confocal microscope RAMOS S120 using 532 nm laser (Ostec Instruments). The photoluminescence spectra and excitation-emission photoluminescence maps were measured by a NanoLog excitation–emission spectrofluorometer (Horiba) with high concentration liquid samples and solid samples placed in frontal mode at 30 degrees.

*Microelectrode Fabrication and Characterization*. Inkjet printing was made using an Autodrop Professional AD-P-8000 with a 50 μm nozzle diameter microdispenser head MD-K- 130 (Microdrop Technologies GmbH; printing parameters summarized in Table S3). Aerosol jet printing has been made using an IDS Nanojet Desktop system. Microelectrodes were printed on 30mm Millicell culture plate inserts (PICM0RG50; Sigma Aldrich / Merck). The insert substrate is made of porous hydrophilic PTFE membrane featuring 0.4 μm pore sizes. For initial biocompatibility testing, 10 parallel lines of 10 mm length incrementally distanced by 1 mm were printed mimicking MEA features.

To fabricate double side printed microelectrode schematics (Figure 1b,c), the membrane substrates were pierced using tungsten micro needles (<1µm tip diameter, 125 µm shaft diameter, World Precision Instruments). This enabled us to produce μm-sized pores that were filled by the inkjet printing with the conductive ink, providing a good conductivity pathway between the two sides of the membrane.

Scanning electron microscopy (SEM) images were obtained with a scanning electron microscope (JEOL JCM-6000PLUS) operating at 15 kV. We used a physical vapor deposition system (Moorfield M307) to deposit a thin (approx. 100 nm) layer of gold on the studied samples, enabling SEM imaging. Surface porosity was evaluated from SEM images using open-sourced FIJI/ImageJ software (calibrating the image under the Analysis → Set Scale function; then with the Image → Adjust → Adjust Threshold function to develop a black/white contrast between the individually resolved pores on the surface; surface porosity was defined as a ratio of the area of the pores obtained using the Analysis → Particle Analysis function vs. total area in the ROI). Electrical impedance spectroscopy was performed in the range from 3 Hz up to 1MHz using Fluke 8846A.

*Organotypic culture*. Cutting solution was prepared by dissolving the following components in distilled water: sucrose 180 mᴍ, KCl 2.5 mᴍ, MgSO_4_ 10 mᴍ, NaH_2_PO_4_ 1.25 mᴍ, NaHCO_3_ 25 mᴍ, glucose 1 mᴍ, ascorbic acid 1 mᴍ, taurine 1 mᴍ, ethyl pyruvate 20 mᴍ, CaCl2 0.5 mᴍ. Artificial cerebrospinal fluid (aCSF) solution was prepared by mixing in distilled water: NaCl 125 mᴍ, KCl 3 mᴍ, MgSO_4_ 1.6 mᴍ, NaH_2_PO_4_ 1.25 mᴍ, NaHCO_3_ 26 mᴍ, glucose 10 mᴍ, CaCl_2_ 2 mᴍ. Osmolarity of both solutions was in the range of 300-310 mOsm L^-1^. Both solutions were sterilized by Stericup Quick Release Filter (Millipore; 0.22um PES Membrane) and carbogenated (95% O_2_/5% CO_2_) for at least 30 minutes before use. ^[3]^ Instruments and beakers were sterilized with autoclave (at 121°C; Prestige Medical Classic Media Autoclave), a microtome, and all the instruments were sprayed with 70% ethanol and allowed for full drying before use. The brains were extracted from rats (postnatal day 7; P7) humanely culled via Animals (Scientific Procedures) Act 1986 (ASPA) Schedule 1 methods; ethical approval was obtained prior the research and all procedures involving animals (rats) have been compliant with the Home Office (UK) guidelines / personal licenses (PP9211565) and the NC3R’s ARRIVE guidelines. The brain was sliced with a thickness of 350 µm in the ice-cold cutting solution. Freshly cut slices were transferred to aCSF continuously bubbled with 95% O_2_/5% CO_2_ and placed on a mesh to rest for 1 hour at room temperature.

Afterwards, the slices were transferred to the organotypic culture media allowing to flush out excess of aCSF. The culture media have been prepared using the following protocol: 50 mL Gibco™ Neurobasal™-A Medium was mixed with 1 mL Gibco™ B-27™ Supplement, 150 µL Gibco™ Gentamicin and 125 µL GlutaMAX™ Supplement.

The slices were placed on the 30mm Millicell culture plate inserts (PICM0RG50; Sigma Aldrich / Merck) with and without microelectrodes (for reference). The inserts with microelectrodes were washed in an aqueous solution of PBS to remove excess PS1 and then sterilized by UV light (1 hour on each side). The inserts were maintained in wells with the culture media at 37°C temperature and 5.0% CO_2_ supply of a culture incubator. The culture media was changed for a fresh one once per 3-4 days to provide a proper nutrient supply.

During the culture period (0-21 days), individual slices were removed from the wells and either fixed for histology or subject to electrophysiological recordings.

Fixed slices were stained for histology by standard H&E (hematoxylin and eosin) staining kit (ab245880; datasheet and protocol - www.abcam.com/ab245880). Brightfield images were digitized from a Toupcam eyepiece camera connected to a binocular microscope (GT Vision) at x40, and cell counts were performed using FIJI.

*Electrophysiological Recordings.* Culture wells containing brain slices as prepared above were transferred to one of two electrophysiology setups to test ensemble activity:

*Glass Microelectrode Rig* consisted of an interface recording chamber (Scientific Systems Design) filled with standard aCSF (see Organotypic Cultures above) and binocular dissecting microscope (Olympus) with dual electrodes connected to a local field potential amplifier (NPI Electronics EXT10-2F) and filter (NPI Electronics LHBF-48X). Glass microelectrodes (1- 3MΩ) filled with aCSF were placed upon recording pads creating a continuous connection ensuring that neither electrode nor pad was damaged. Ensemble activity was recorded at a total amplification of 1000x, with low and high pass filters of 700 Hz and 0.5 Hz, respectively. Signals were digitized at 2 kHz using a CED 1401 connected to a Windows PC running Spike2 version 8.

*Solid Electrode Rig* was based on a Kerr Scientific In Vitro Brain Slice System, with the culture well placed in the central chamber and filled with aCSF. Insulated solid wire electrodes (Kerr Scientific) were placed onto the recording pads firmly, creating a slight deformation in the PTFE substrate to ensure maximal connection. Signals were amplified 250x with the Kerr isolated amplifier, AC-coupled at 0.5 Hz and low-pass filtered at 500 Hz, digitized at 2 kHz using a CED 1401 connected to a Windows PC running Spike2 version 8. In all cases, cultured slices were perfused with a minimal amount of aCSF bubbled with 95% O_2_/5% CO_2_ and left in the chamber to equilibrate for 15 minutes. Locations of printed electrode pads were determined under the microscope, and the pad(s) closest to the cell body layer of the hippocampus, or to layers 2/3 of the neocortex, were chosen. Multi-unit activity was elicited after a 10-minute baseline period via bath addition of 20 µM gabazine (GABAa) to disinhibit the slices. Electrodes were removed from pads and placed into the aCSF after recording to ensure that multi-unit activity was recorded via the contact pads rather than transmitted through the fluid.

## Acknowledgements

P.L. and S.D.G. acknowledge support of the British Academy, Royal Academy of Engineering and Royal Society (Academies Partnership in Supporting Excellence in Cross-disciplinary research award - APEX award, AA21\100133 APEX Awards 2022). Dr. Emily Allwright and Dr. Neil Chilton are acknowledged for technical assistance in aerosol jet printing using the IDS Nanojet system at Printed Electronics Ltd.

## Data Availability Statement

The data that support the findings of this study are available from the corresponding author(s) upon reasonable request.

Received: ((will be filled in by the editorial staff))

Revised: ((will be filled in by the editorial staff))

Published online: ((will be filled in by the editorial staff))

## Funding

British Academy, Royal Academy of Engineering and Royal Society (Academies Partnership in Supporting Excellence in Cross-disciplinary research award - APEX award, AA21\100133 APEX Awards 2022).

## Supporting Information

Supporting Information is available from the Wiley Online Library or from the author.

Table of Contents figure (110 mm × 20 mm)

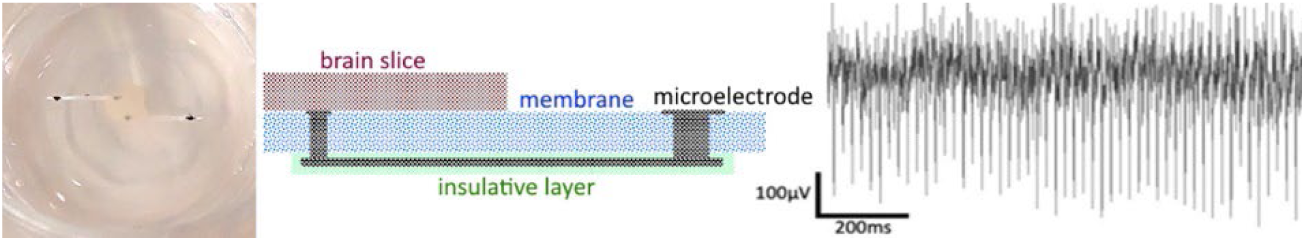

Table of Contents text:

The highly porous microelectrodes have been designed and printed on culture membranes allowing to record electrophysiological neural activity for rodent brain slices. To keep the biocompatible nanoporous structure, the microelectrodes and insulative layer were fabricated on the bottom of culture membranes with only small connector pads added on the top.

## Supporting Information

### **1.** Ink characterization

We measured surface tension (*γ*), density (*ρ*), and dynamic viscosity (*η*) of produced inks to analyze their inverse Ohnesorge number 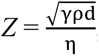, where *d* = 50 µm is nozzle diameter in our inkjet printer (Table S1).

**Table S1.**
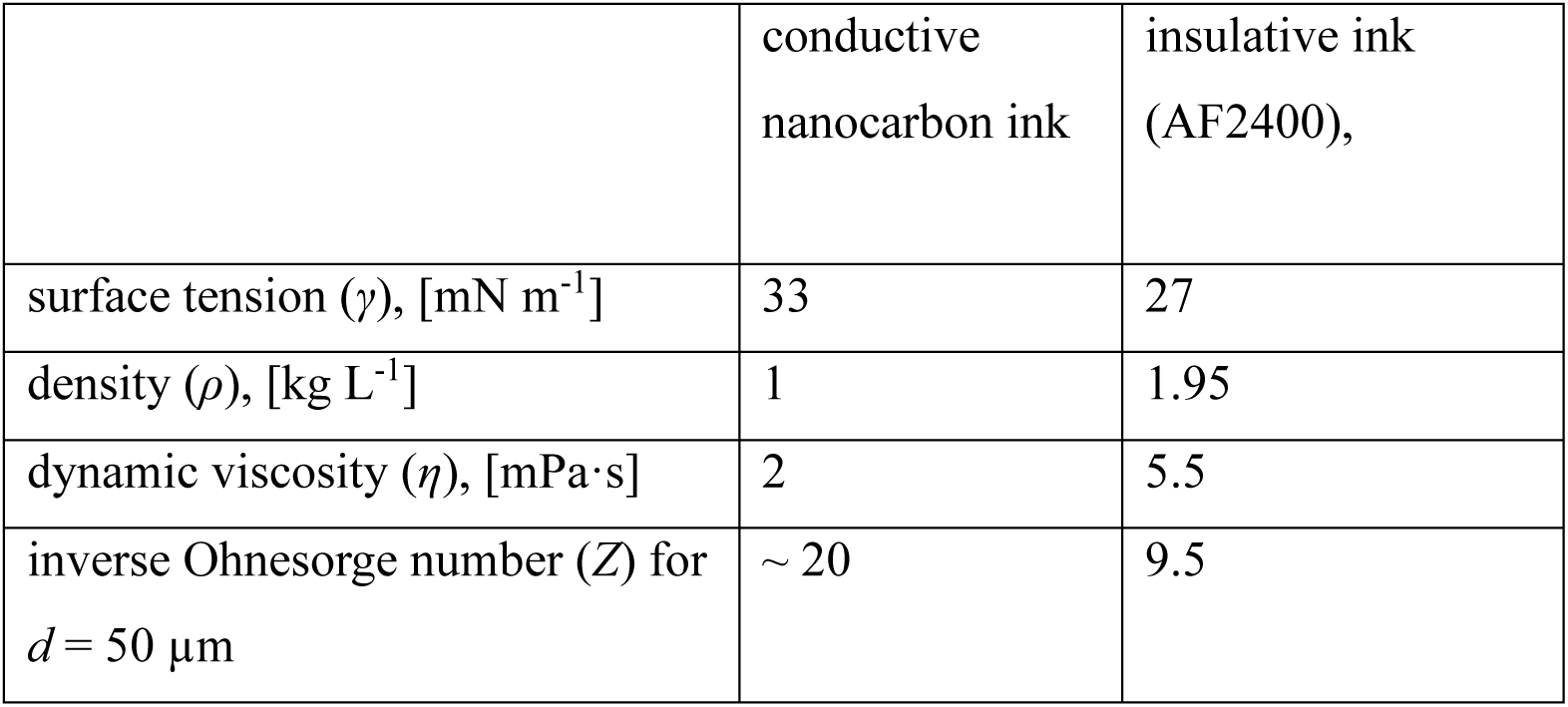
Ink formulation parameters.

|  | conductive<br>nanocarbon ink | insulative ink<br>(AF2400), |
| --- | --- | --- |
| surface tension ($\gamma$), [$\text{mN m}^{-1}$] | 33 | 27 |
| density ($\rho$), [$\text{kg L}^{-1}$] | 1 | 1.95 |
| dynamic viscosity ($\eta$), [$\text{mPa}\cdot\text{s}$] | 2 | 5.5 |
| inverse Ohnesorge number ($Z$) for<br>$d = 50\ \mu\text{m}$ | $\sim 20$ | 9.5 |

**Figure S1.**
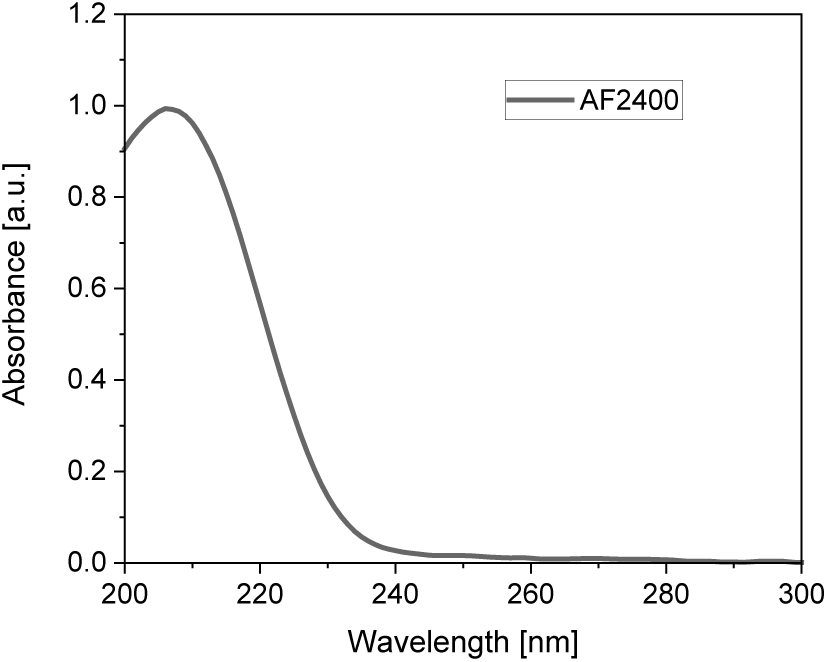
Absorption spectra of perfluorodecalin solution of AF2400 (5 g L^-1^; 10 mm quartz cell).

### **2.** Additional SEM characterization of PTFE membrane

**Figure S2.**
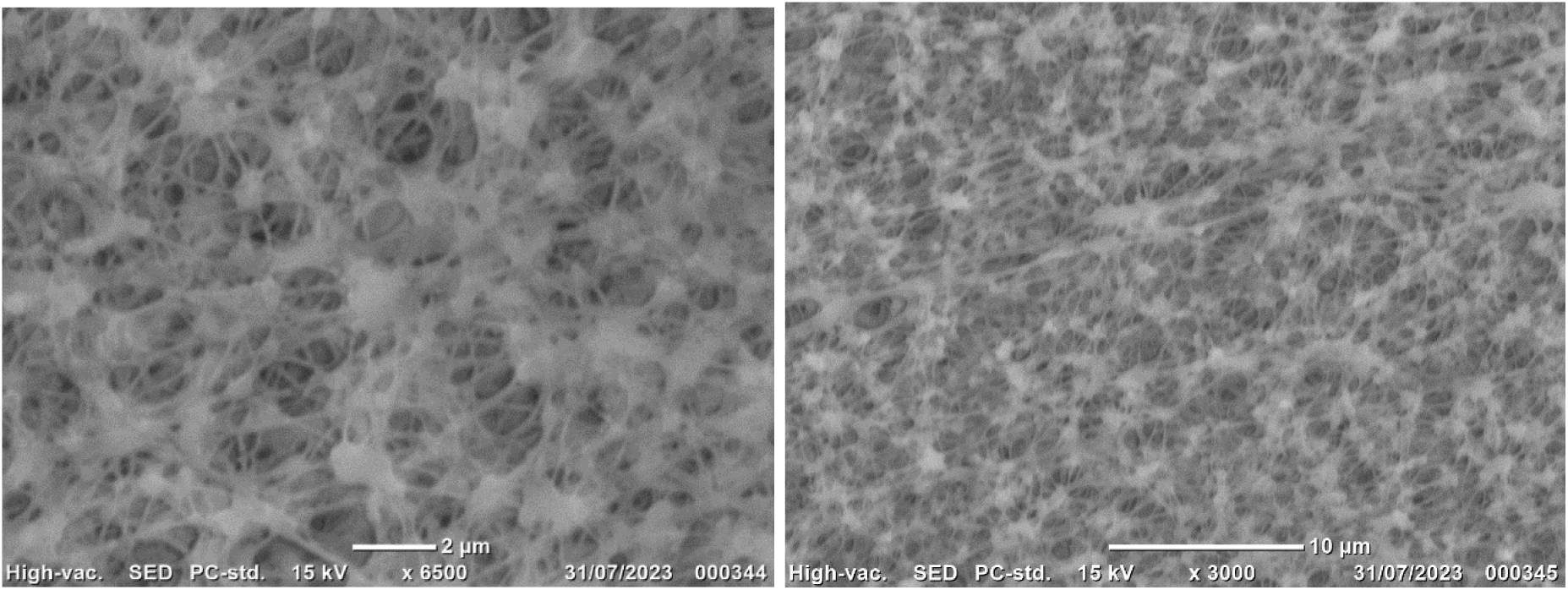
SEM images for neat PTFE membrane of culture insert substrate

### **3.** Contact angle characterization

Contact angles (*θ*) were measured for nanocarbon (Figure S3) and AF2400 (Figure S4) inks on quartz and PTFE membranes.

**Figure S3.**
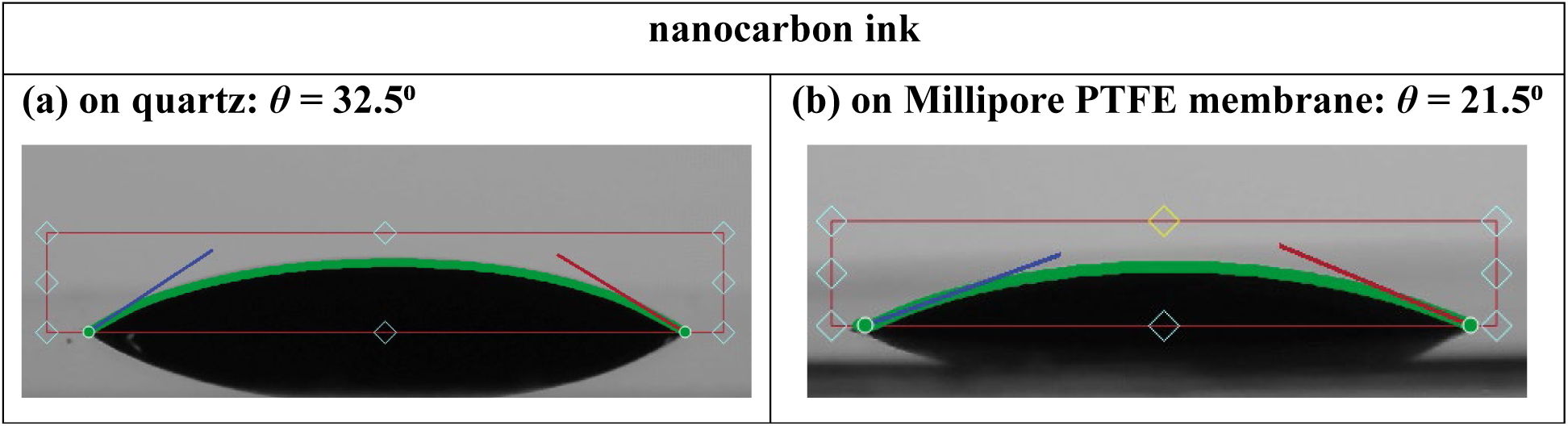
Images of nanocarbon ink drops for contact angle measurements on (a) quartz and (b) the Millipore PTFE membrane.

It should be highlighted that the nanocarbon ink (based on water/propylene glycol solvent) drops are slowly drying on the PTFE membranes due to very gradual wetting of nanoporous surface.

AF2400 ink (based on perfluorodecalin as solvent) demonstrates very quick wetting of the membrane surface within 1^st^ second after drop lands the surface. Such behavior features dynamic contact angle measurements (Figure S4 and Figure S5). Besides, due to quick wetting of AF2400 ink, the inkjet printed lines are much wider than the nanocarbon lines. The video recording for results presented in Figure S4 and S5 is available as part of the Supporting Information.

**Figure S4.**
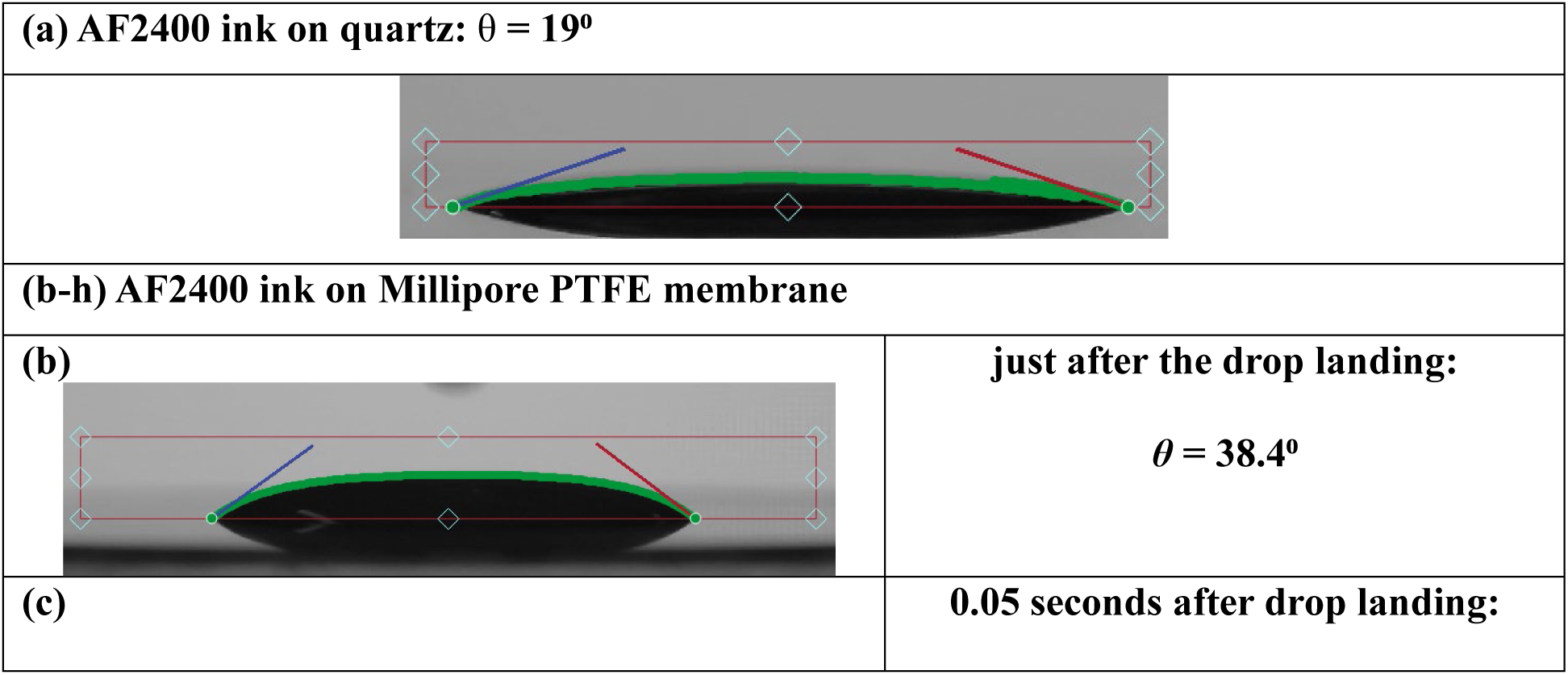
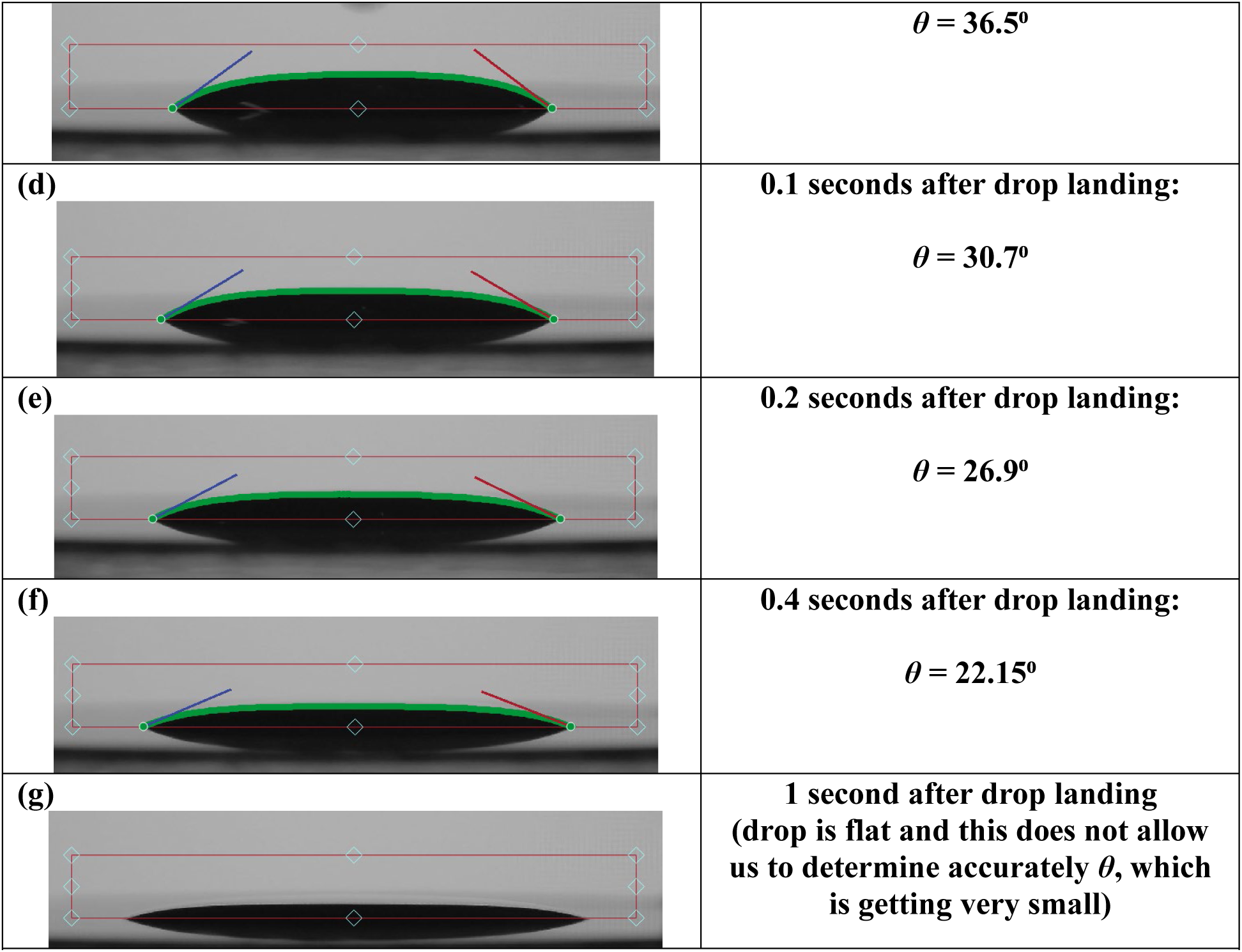
(a-g) Images of AF2400 ink drops for contact angle measurements on quartz and (b-g) the Millipore PTFE membrane.

**Figure S5.**
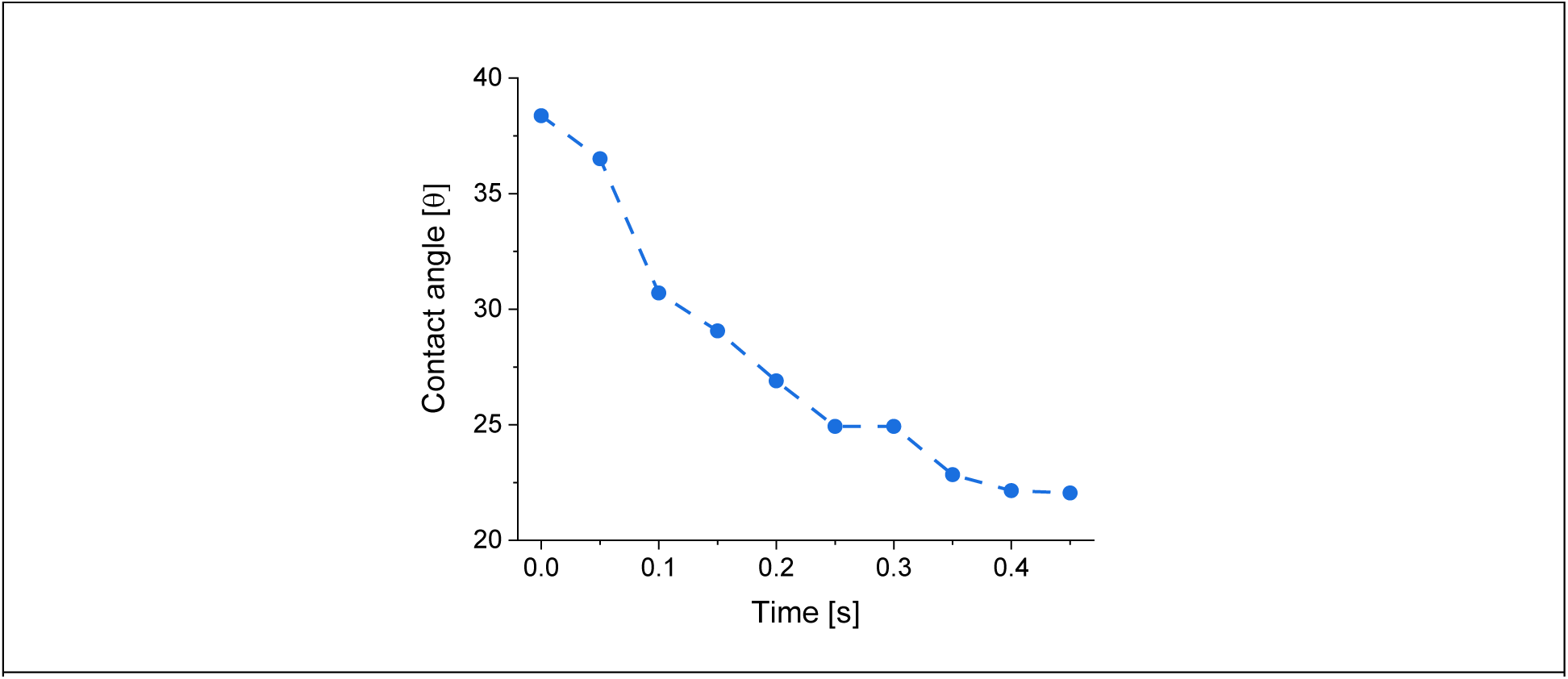
Temporal changes of AF2400 ink contact angle on the Millipore PTFE membrane.

### 4. Additional photoluminescence characterization

**Figure S6.**
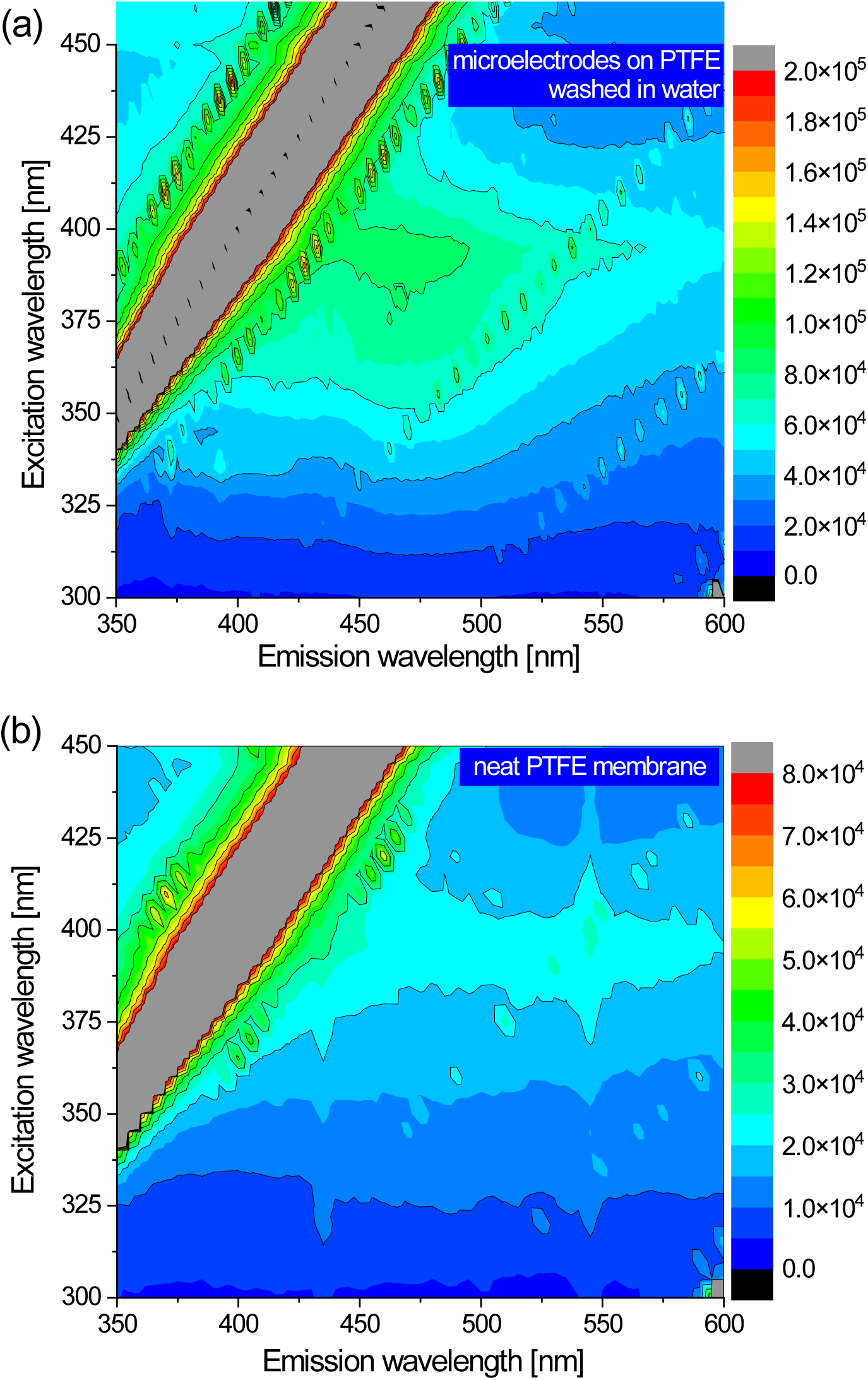
Excitation-emission photoluminescence (PL) map of the (a) inkjet printed microelectrodes on PTFE membranes after washing in water and (b) neat PTFE membranes.

### **5.** Parameters of the Constant Phase Element (CPE) equivalent circuit used to model the impedance spectra

The impedance spectra in Figure 5a were fitted by the equivalent circuit in Figure 5b.

The circuit includes the CPE that represents the capacitive behavior of the microelectrodes, and impedance of CPE (Z_CPE_) is defined as following: ^[1,2]^

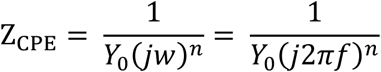

where ω is angular frequency, Y_0_ and n are the characteristic parameters of the CPE, with n changing in the range of 0 to 1. Evaluation of Y_0_ allows us to calculate the double layer capacitance (C_dl_) of the microelectrode:

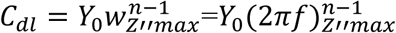

where 𝑗𝑗_𝑍𝑍′′*max*_ is the angular frequency having the imaginary part of impedance at its maximum.

**Table S2.**
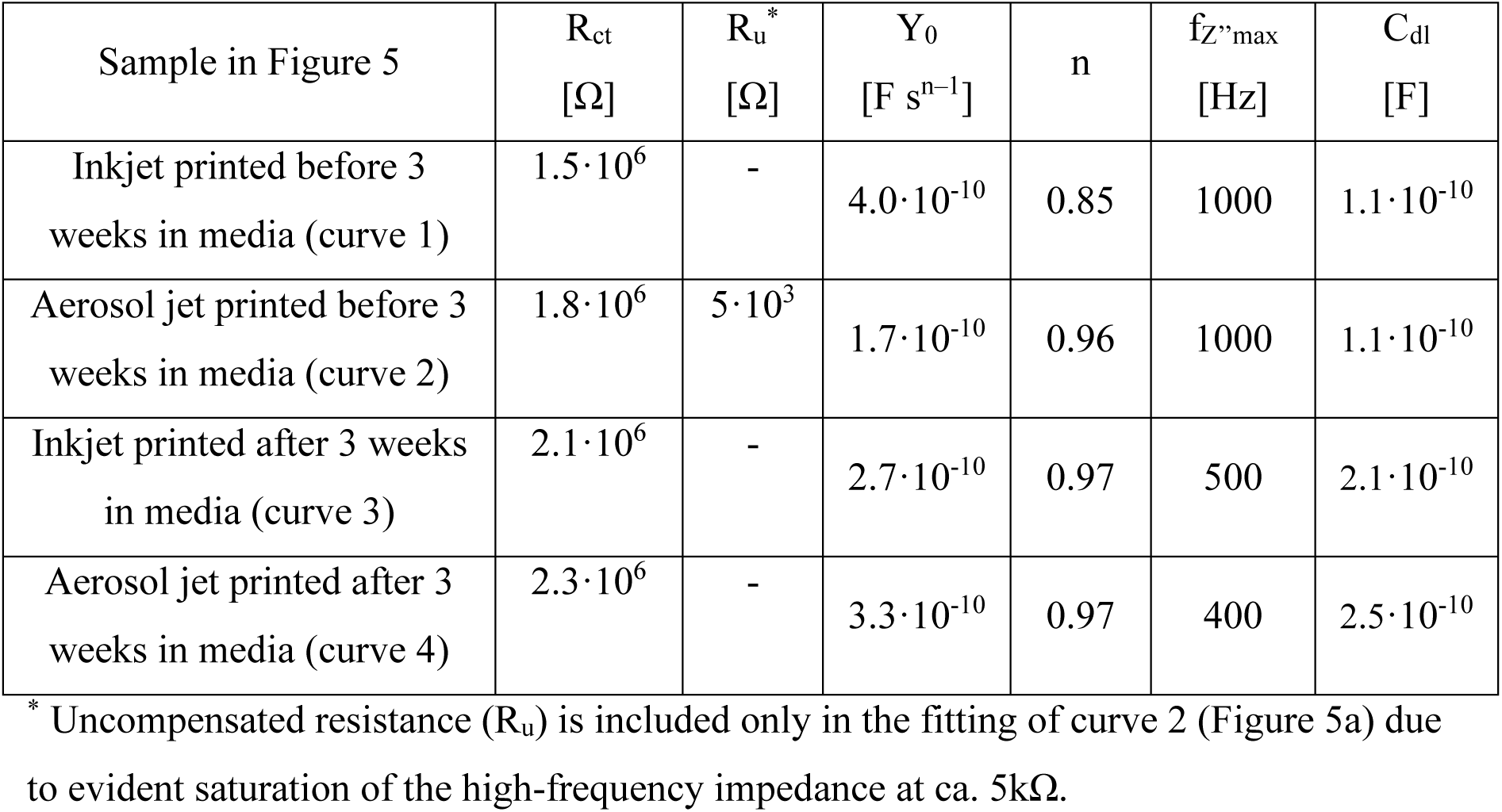
Parameters of the equivalent circuit in. **Figure 5b used to fit the impedance spectra in Figure 5a**

The effective surface area (ESA), also known as the electrochemically active surface area, is the actual area of an electrode available for electrochemical reactions. It differs from the geometric surface area, which is the physical, two-dimensional size of the electrode. The ESA accounts for the micro- and nanoscopic roughness, porosity, and surface chemistry that create a much larger actual surface. ESA can be determined from the C_dl_ using the relationship:

ESA = C_dl_ / C_s_. The specific capacitance (C_s_) is a known constant for the electrode material, and considering 20 μF·cm^−2^ for a perfectly smooth surface of an electrode, ^[3]^ the ESA of the studied microelectrodes has a range of 10^-5^ cm^2^.

### **6.** Frequency response for noise level

**Figure S6.**
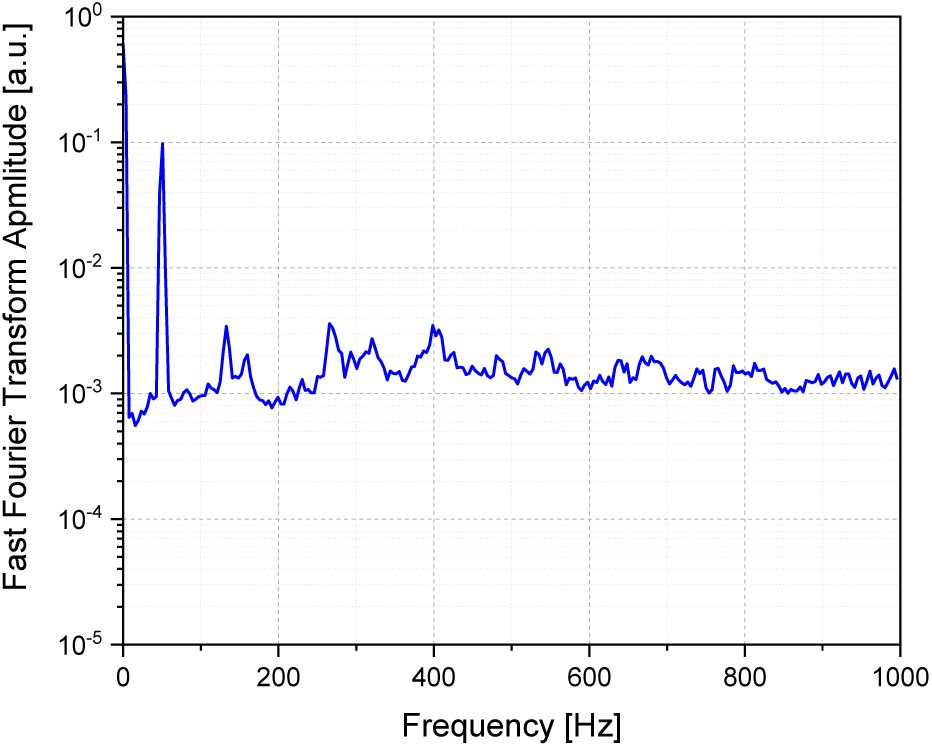
Fast Fourier Transform (FFT) spectrum of the noise showing no abnormalities apart of 50 Hz spike.

### **7.** Printing Parameters

**Table S3.** Printing parameters for inkjet printing process*.

|  | conductive<br>nanocarbon ink | insulative ink<br>(AF2400), |
| --- | --- | --- |
| Printing (dispenser) head voltage range, V | 60 – 126 | 50 – 85 |
| Printing (dispenser) head pulse width, μs | 25 – 50 | 38 – 50 |
| Drop frequency in ‘Continuous Mode’, Hz | 200 | 200 |
| Holding pressure range, mbar | (-5) – (-11) | (-12) – (-14) |
| Dispensing step for line printing, μm | 50 | 70 |
| Number of printing passes | 10 | 20 |
| Average width of the printed line, μm<br>(at above number of passes) | 100 | 200 |
| Minimum width of the printed line, μm<br>(at above number of passes) | 90 | 160 |

\* All printing has been performed at room temperature (no dispenser head heating) in cleanroom environment.

## Notes

### Competing Interest Statement

The authors have declared no competing interest.

### Summary of Updates

Results and Discussion section is updated, adding extra characterisation and discussion on the ink and the microelectrode characterization; one more co-author added; Supporting information file is combined with the manuscript following the template for the revised version.

